# A cryo-EM processing pipeline for microtubules using CryoSPARC

**DOI:** 10.64898/2026.02.24.703950

**Authors:** Daniel Zhang, Hugo Muñoz-Hernández, Pavel Filipcik, Kushal Sejwal, Yixin Xu, Sung Ryul Choi, Michel O. Steinmetz, Michal Wieczorek

**Author notes:** These authors contributed equally (Hugo Muñoz-Hernández, Pavel Filipcik). Medical Sciences Doctoral Training Centre, University of Oxford, Oxford, UK. AAX Biotech AB, Nanna Svartz väg 4, 171 65 Solna, Sweden.

## Abstract

Microtubules are cytoskeletal filaments typically characterized by a discontinuous helical lattice of α/β-tubulin heterodimers. Microtubules can also adopt variable lattice architectures both *in vitro* and in cellular contexts. Pseudo-helical averaging processing strategies have been developed to generate cryo-EM reconstructions of microtubules with and without decorating protein-binding partners, but these pipelines can be difficult to implement for the average user, especially for undecorated filaments. Here, we describe MiCSPARC, a cryo-EM processing pipeline developed around CryoSPARC (Punjani et al., 2017), which leverages automated particle picking and fast 3D refinement times in CryoSPARC to determine structures of both decorated and undecorated microtubules. We generated reconstructions of undecorated GDP microtubules, as well as kinesin-1 motor domain-decorated GMPCPP filaments at resolutions of up to 2.8 Å, demonstrating the robustness of the pipeline. Based on its convenient implementation and ability to routinely generate high-resolution, seam-corrected microtubule reconstructions, MiCSPARC should provide a valuable tool for understanding microtubule dynamics, microtubule-associated proteins, and microtubule-targeting agents.

## Introduction

Microtubules are cytoskeletal polymers built from α/β-tubulin heterodimers, which self-associate in a head-to-tail manner into linear, polarized protofilaments; β-tubulin, a site of GTP binding and hydrolysis, points towards the microtubule’s fast-polymerizing or “plus” end, while α-tubulin, containing a non-exchangeable GTP binding site, points towards the slow-polymerizing or “minus” end [reviewed in (Brouhard and Rice, 2014; Akhmanova and Steinmetz, 2015)]. In canonical microtubules, 13 α/β-tubulin protofilaments associate laterally in the A-type configuration, resulting in a hollow cylindrical structure and a “seam” that forms between only two protofilaments marked by heterotypic B-type α- and β-tubulin lateral contacts (Wade et al., 1990; Sui and Downing, 2010). The seam therefore complicates helical averaging in cryo-electron microscopy (cryo-EM) single particle analysis (SPA) because the α- and β-tubulin monomers are structurally similar and difficult to distinguish from one another during classifications. Introducing a decorating protein that forms a 1:1 complex with each α/β-tubulin heterodimer can help identify the seam (Zhang and Nogales, 2015), but can also physically distort the microtubule lattice (Lacey et al., 2019; Zhang et al., 2018). Moreover, differences in microtubule lattice architectures, including, e.g., variable protofilament numbers, tubulin dimer spacing along the lattice, supertwists along the filament, as well as partial binding occupancy of associated proteins to *in vitro* polymerized microtubules, all further limit achievable resolutions and the interpretability of reconstructions generated with conventional processing pipelines.

Several strategies have been developed to determine structures of decorated microtubules (Cook et al., 2020; Zhang and Nogales, 2015; Debs et al., 2020). However, these methods experience limitations that make routine and fast microtubule reconstructions difficult to achieve for average cryo-EM users. For example, accurate 3D references containing correctly positioned decorators relative to α/β-tubulin are crucial for initial alignments, but current methods use either existing structures as references based on loose structural similarity (Cook et al., 2020), or else generate references entirely *in silico* (Zhang and Nogales, 2015), both of which can introduce potential sources of bias. Particle pre-averaging is also used to increase signal-to-noise for initial alignments (Cook et al., 2020; Zhang and Nogales, 2015), but adds unnecessary complexity, since the so-called “super-particles” need to first be well aligned in 2D to start being truly helpful in subsequent processing steps. Lastly, existing strategies don’t yet take advantage of the accessibility and faster refinement times in software packages like CryoSPARC (Punjani et al., 2017). As a result, it remains difficult and time consuming to determine cryo-EM structures of decorated microtubules, and seam-corrected, undecorated filament reconstructions remain out of reach for most users, limiting potential progress in the field of microtubule structural biology.

Here we describe MiCSPARC, a microtubule processing pipeline developed around CryoSPARC that leverages improved automated particle picking, protofilament number sorting, seam searching, and fast 3D refinement times in CryoSPARC to accurately reconstruct both decorated and undecorated microtubules. We provide example MiCSPARC reconstructions of seam-corrected microtubules, including a seam-corrected, undecorated filament at overall ∼3.6 Å resolution, with α/β-tubulin in a single protofilament reconstruction reaching 3.0 Å. The ability of MiCSPARC to reliably generate high-quality microtubule reconstructions should accelerate structural studies of undecorated microtubules and filaments in complex with microtubule-associated proteins or microtubule targeting agents.

## Results

### Developing MiCSPARC to reconstruct a kinesin-1 motor domain-decorated, GMPCPP microtubule

We developed MiCSPARC (Appendix 1) using a test dataset of guanosine-5’-[(α,β)-methyleno] triphosphate (GMPCPP)-stabilized microtubules decorated with the microtubule-binding motor domain of KIF5A (Supplementary Table 1), a member of the kinesin-1 protein family. Microtubules decorated with kinesin motor domains are typically more easy to align and average than undecorated filaments (Alushin et al., 2014), and the relatively large kinesin motor domain acts as a fiducial that helps localize particle positions relative to the seam in downstream processing steps (Ti et al., 2018).

We first optimized the particle picking step, which normally involves tracing the microtubule and extracting overlapping segments every ∼8 nm as 2D particles. Manual selection of microtubule segment start-and-end points, as previously described (Cook et al., 2020), is a time-consuming, tedious process that does not easily allow for curved filament picking and can be prone to user bias. CryoSPARC’s filament tracer reliably picks microtubules without significant user intervention, but it also tends to incorrectly assign particles from different filaments into the same filament. Moreover, it can miss portions of filaments entirely, particularly near the edges of micrographs or in densely populated regions.

To address these challenges, we developed a script that extrapolates coordinates from the Filament Tracer tool in CryoSPARC to more completely select and keep track of microtubule filaments. The script incorporates both linear and quadratic curve fitting to account for slight, naturally occurring bends in microtubules imaged by cryo-EM. By iteratively comparing best fit models of each filament to particles picked across all filaments, proximal particles are dynamically reassigned, mitigating the effects of incorrect assignments from CryoSPARC’s filament tracer tool. In addition to generating typically larger numbers of starting particles, we found that a robust automated picking procedure also alleviates the need to generate locally averaged “super-particles”, which is presently required by other methods for accurate initial particle alignments (Zhang and Nogales, 2015; Cook et al., 2020).

Next, we focused on the challenge of generating 3D references for supervised classification. In current microtubule reconstruction pipelines, 3D references are either used directly from “standard” datasets of undecorated and decorated microtubules [e.g., (Sui and Downing, 2010; Cook et al., 2020)], or else synthetic “decorated” references mimicking the addition of the binding protein needed to be generated (Zhang and Nogales, 2015), which requires accurate knowledge of the decorator’s structure and binding mode on the tubulin lattice. In our hands, synthetic or a priori references can work when the shape and location of the binder are already known but will tend to fail without a priori knowledge. Even with such knowledge, slight mismatches between reference and real structures due to, e.g., unexpected helical supertwists (Zhang et al., 2015, 2018) lead to non-converging 3D refinements.

In MiCSPARC, we dealt with these challenges by reconstructing a single, roughly aligned protofilament from the data - which was generated from a set of initial, reference-free helical refinements of all picked particles - to generate a set of synthetic references with any desired theoretical helical parameters (Figure 1A). These references were then used for supervised heterogeneous refinement in CryoSPARC. This resulted in a robust sorting of microtubule architectures (Figure 1B), as the dominant microtubule type in the kinesin-decorated dataset was 14 protofilaments with a 3-start helix, as expected for microtubules polymerized in the presence of GMPCPP (Hyman et al., 1992). Using information from this supervised classification, MiCSPARC then employs a smoothing algorithm to assign a distinct protofilament number to each picked microtubule (Cook et al., 2020), which is also needed for generating accurate seam-corrected microtubule reconstructions below.

**Figure 1.**
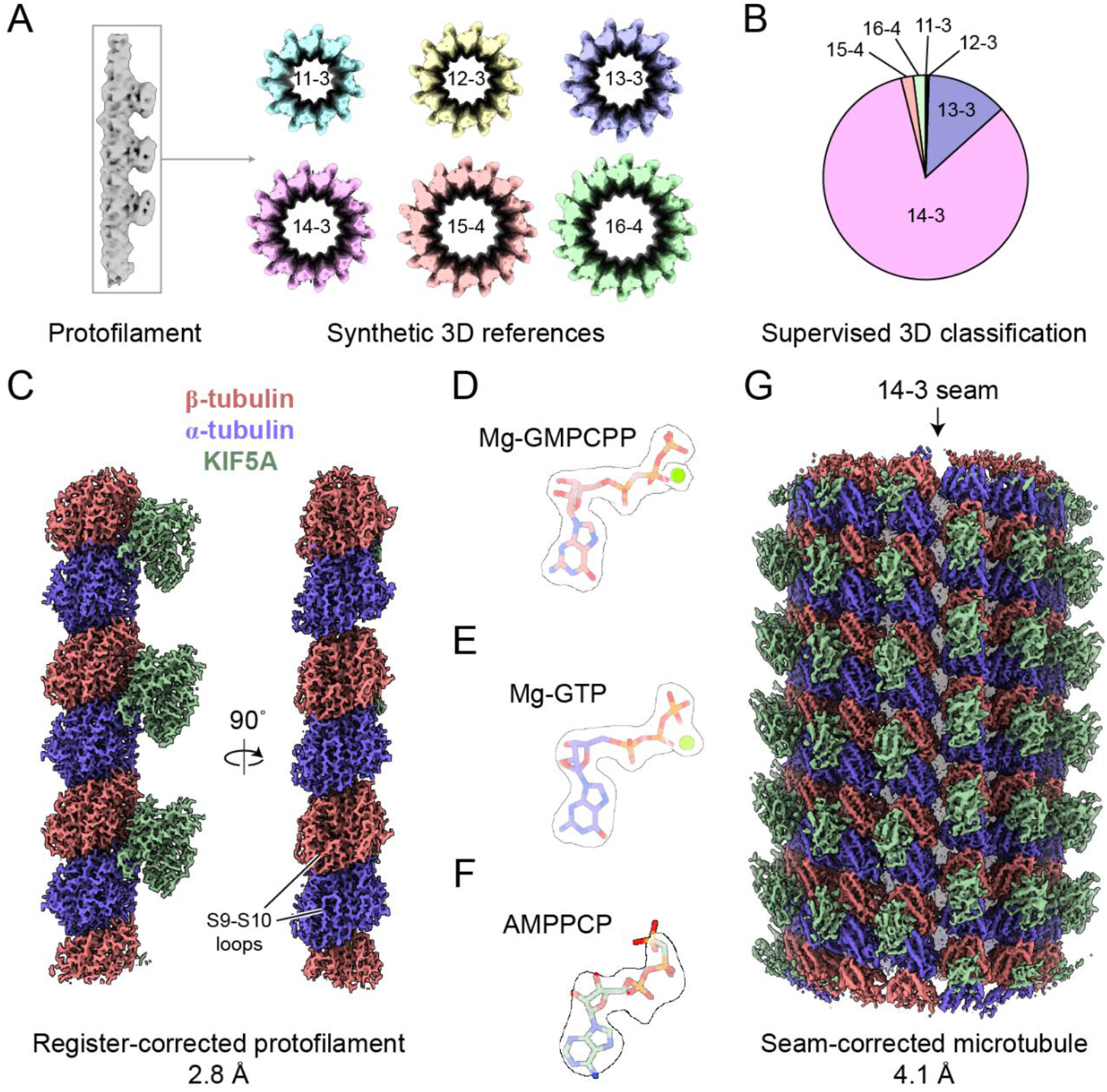
MiCSPARC reconstruction of a KIF5A motor domain-decorated, GMPCPP microtubule. **A)** Initial rough signal-subtracted reconstruction of a single KIF5A-decorated protofilament (left), which was used to then computationally generate synthetic 3D references for supervised 3D classification (right; coloured reconstructions; protofilament number and helical “start” numbers are labeled). **B)** Resulting distribution from supervised 3D classification and 3D class “smoothing” of the KIF5A-decorated microtubule dataset using the references in A). **C)** Two views of a final MiCSPARC refinement of a single KIF5A-decorated protofilament after symmetry expansion and α/β-tubulin register correction. Obvious differences in α-vs β-tubulin S9-S10 loops are indicated. The map was postprocessed using EMReady (He et al., 2023). **D)** Mg-GMPCPP in β-tubulin (stick representation) in the corresponding density (transparent surface representation). **E)** Mg-GTP in α-tubulin (stick representation) in the corresponding density (transparent surface representation). **F)** AMPPCP in KIF5A (stick representation) in the corresponding density (transparent surface representation). Densities in D)-F) are thresholded at the same levels. **G)** Seam-view of the seam-corrected, 14 protofilament and 3-start microtubule reconstruction obtained using MiCSPARC. The location of the seam is indicated by an arrow.

The subsequent processing steps in MiCSPARC have two main goals: i) protofilament reconstruction, which allows reaching higher resolutions locally but also unifies incorrectly assigned Euler angles from global refinements; and ii) generating a seam-corrected microtubule reconstruction, which uses the alignment information determined in i). To reconstruct a protofilament (goal (i)), we made the commonly held assumption that each microtubule within a micrograph does not change its lattice structure. Microtubule particles from a given protofilament number from the 3D classification step were first refined, and per-microtubule in-plane (psi) rotation angles were then unified, following previously described methods (Cook et al., 2020). Psi-unified particles were re-imported and locally refined in CryoSPARC. Angles representing rotations around the filament axis (phi) were then also iteratively unified on a per-microtubule basis, using a modification of previous methods (Cook et al., 2020) (see Methods).

After a final C1 refinement using the updated priors, each particle within a given 3D class was next symmetry expanded using the previous helical refinement outputs, and a single protofilament was masked and again subjected to a local refinement. 3D classification without alignment in CryoSPARC was employed to identify two protofilament maps that are translated by one tubulin monomer relative to one another; this is identifiable by the shifted presence of kinesin motor domain decorator. These maps were used as references for supervised 3D classification of all symmetry-expanded protofilament particles. Alignment of one of the resulting classes to the other followed by final refinement steps resulted in a 2.8 Å reconstruction of the protofilament, in which all particles contain the correct α/β-tubulin register (Figure 1XFigure 1C, Supplementary Figure 1A-B and Supplementary Table 2). The quality of the reconstruction was also sufficient to identify all expected nucleotide states in the kinesin motor domain, α-tubulin, and β-tubulin, including coordinated magnesium ions representing pre-hydrolysis states (Figure 1D-E, Supplementary Figure 2A and Supplementary Table 3).

To reconstruct the seam-corrected, decorated microtubule (goal (ii)), the sets of particles classified during α/β-tubulin register correction above were used to determine the position of the microtubule seam. In principle, the seam is the position at which the protofilament register changes. Thus, the position of the seam can be determined where consecutive symmetry-expanded and well-aligned particles switch from occupying one register class to the other within the same microtubule filament. However, as the 3D classification during α/β-tubulin register correction is not perfect, corrective measures can also be applied to improve the confidence of the assignment. We therefore scored each position on the likelihood that it is the seam position, based on the registers of neighboring protofilaments. The score increases if the protofilaments on one side of the position have been assigned to the same register, and a different register to the protofilaments on the opposite side. Due to the previous phi angle correction, the order of protofilaments in each image along the microtubule can be assumed to be the same. Thus, the seam likelihood scores for each position can be averaged along the microtubule to give a more robust estimation of the most likely seam position. The particles at the most likely seam position are then returned to be extracted and refined as the seam-corrected microtubule.

For the KIF5A-decorated dataset, this procedure generated a 4.2 Å resolution reconstruction of the decorated microtubule (Figure 1G, Supplementary Figure 1C-D and Supplementary Table 2), in which the seam of the 14 protofilament, 3-start microtubule can be identified by an obvious shift in KIF5A decorating densities as well as differences in the S9-S10 loops of α- and β-tubulin. Together, these results demonstrate the effectiveness of MiCSPARC in generating high-resolution 3D reconstructions of α/β-tubulin bound to decorating fragments, at the level of both the protofilament and the complete microtubule polymer.

### MiCSPARC reconstruction of an undecorated GDP microtubule

Microtubule-binding proteins typically recognize a feature of the α/β-tubulin heterodimer. For example, the kinesin-1 motor domain binds specifically to the interface between α- and β-tubulin within the tubulin heterodimer. This facilitates identification of the α/β-tubulin register along each protofilament and therefore correcting for the seam, which is why kinesin motor domain decoration has been used extensively in many previous microtubule structural studies (Alushin et al., 2014; Ti et al., 2018). However, it has also been reported that certain microtubule-binding domains, including from kinesins, can induce structural changes in the tubulin lattice (Lacey et al., 2019; Zhang et al., 2018), potentially confounding interpretation of tubulin properties from high-resolution decorated microtubule structures.

We therefore next asked whether MiCSPARC could be used to classify and seam-reconstruct a microtubule lacking any binding proteins. This presents a more difficult processing problem, due to the necessity of the subclassification of subtle differences between α- and β-tubulin, such as the S9-S10 loops, in two proteins with otherwise highly similar folds (Nogales et al., 1995). We collected a cryo-EM dataset of microtubules polymerized in the presence of GTP (Supplementary Table 1), creating mainly GDP-containing lattices (Alushin et al., 2014), and used MiCSPARC’s automated filament tracing, reference generation, and supervised classification pipelines to classify the microtubules into distinct lattice architectures. Consistent with previous findings, MiCSPARC sorted our spontaneously polymerized microtubule images into predominantly 14-3 and 13-3-protofilament architectures (Figure 2A-B). Gratifyingly, we were next able to use MiCSPARC to distinguish between α- and β-tubulin and generate a reconstruction of an α/β-tubulin protofilament at 3.0 Å resolution (Figure 2C, Supplementary Figure 3A-B and Supplementary Table 2). Consistent with current models (Alushin et al., 2014; Hyman et al., 1992), the resulting map shows that α/β-tubulin is in a “compacted” microtubule lattice-bound conformation. Further, at these resolutions, nucleotide features can be clearly distinguished, revealing a magnesium ion and GTP at the nucleotide binding site of α-tubulin, while β-tubulin exhibits only GDP (Figure 2D-E and Supplementary Figure 2B), consistent with biochemical studies of α/β-tubulin (Menéndez et al., 1998).

**Figure 2.**
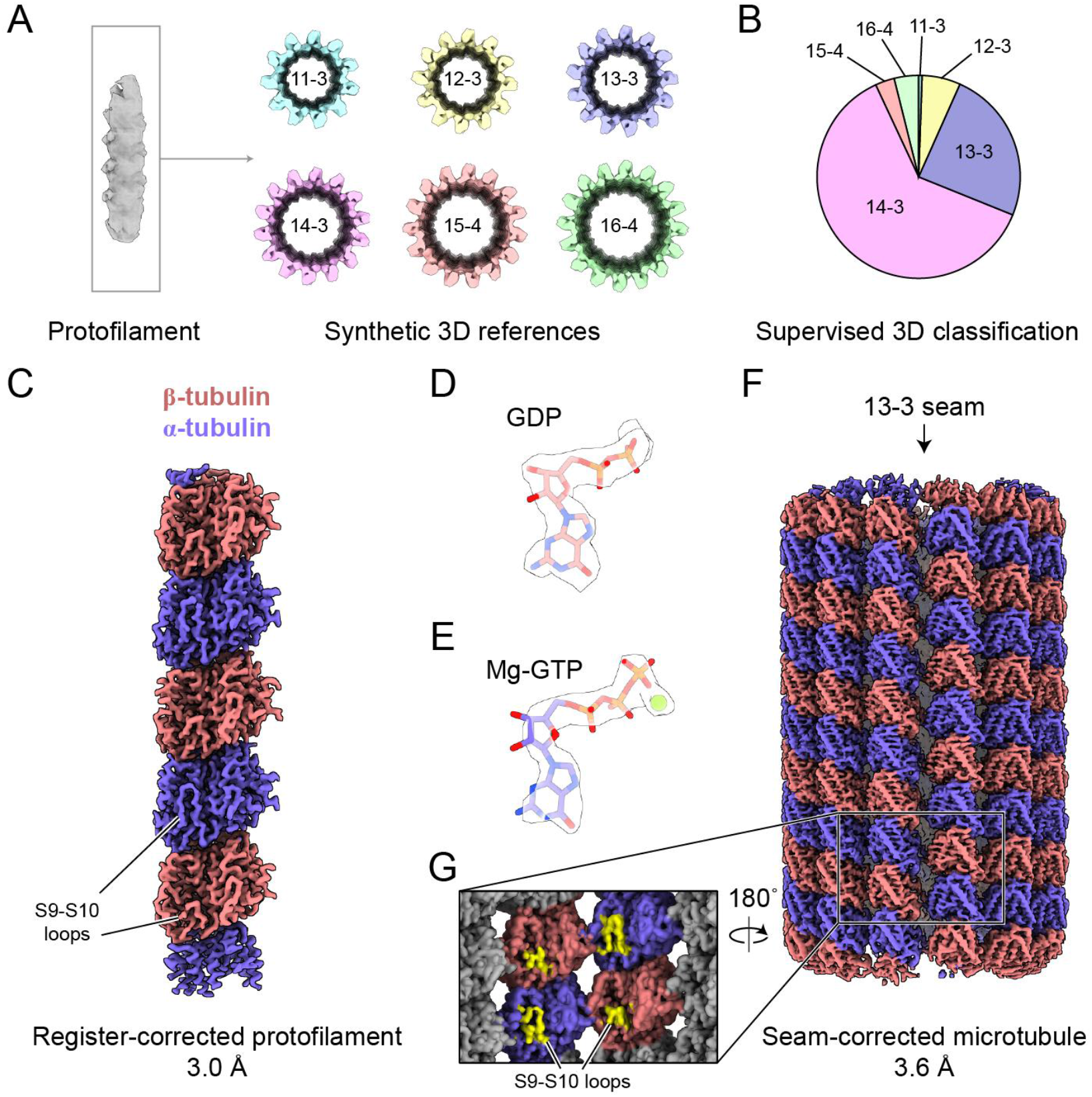
MiCSPARC reconstruction of an undecorated, dynamic microtubule. **A)** Initial rough signal-subtracted reconstruction of a single protofilament (left), which was used to then computationally generate synthetic 3D references for supervised 3D classification (right; coloured reconstructions; protofilament number and helical “start” numbers are labeled). **B)** Resulting distribution from supervised 3D classification and 3D class “smoothing” of the undecorated microtubule dataset using the references in A). **C)** Lumenal view of a final MiCSPARC refinement of a single undecorated protofilament after symmetry expansion and α/β-tubulin register correction. Obvious differences in α-vs β-tubulin S9-S10 loops are indicated. The map was postprocessed using EMReady (He et al., 2023). **D)** GDP in β-tubulin (stick representation) in the corresponding density (transparent surface representation). **E)** Mg-GTP in α-tubulin (stick representation) in the corresponding density (transparent surface representation). **F)** Seam-view of the seam-corrected, 13 protofilament and 3-start microtubule reconstruction obtained using MiCSPARC. The location of the seam is indicated by an arrow. **G)** Zoomed-in and rotated view of the seam of the microtubule reconstruction in F). Differences in α-vs β-tubulin S9-S10 loops across the seam are indicated.

Re-extracting these aligned particles to reconstruct the microtubule (3.5 Å; Figure 2F-G, Supplementary Figure 3C-D and Supplementary Table 2) revealed well-defined S9-S10 loops in α- and β-tubulin and heterotypic contacts at the seam. This suggests that MiCSPARC accurately detects the seam of even undecorated microtubules, and therefore it accurately assigns the phi angles in the protofilament reconstruction step via a possibly more robust algorithm compared with previous approaches (see Methods) (Cook et al., 2020). Altogether, these results demonstrate that MiCSPARC can reliably generate high-resolution, seam-corrected reconstructions of not only decorated microtubules, but also more challenging undecorated microtubule specimens.

## Discussion

We have described MiCSPARC, a cryo-EM processing pipeline for microtubules that leverages the CryoSPARC SPA platform. The key advantages of MiCSPARC over previous methods include: (1) automated particle picking; (2) improved a priori 3D reference generation for supervised 3D classification of different microtubule architecture types; (3) more robust reconstructions of undecorated or sparsely decorated microtubules with correctly-estimated seam positions; and (4) enhancements in processing speed due to CryoSPARC’s faster refinement times compared to alternatives, its user-friendly interface, and its job queueing system, all of which enhance throughput. While MiCSPARC serves as a reliable starting point for reconstructing microtubules from cryo-EM images, achieving optimal reconstructions may still require additional refinement steps either in CryoSPARC itself, or with other established cryo-EM SPA software packages like RELION (Scheres, 2012).

MiCSPARC has several areas for improvement in future versions. These could include further optimization of automated particle picking, 3D classification during microtubule architecture sorting, as well as α/β-tubulin register correction, and seam correction. Moreover, challenges persist in handling decorated filaments exhibiting partial or flexible binding to the microtubule lattice, and the 3D reference generation process still requires users to suggest microtubule architecture types, which may lead to misclassification of unusual helical supertwists or lattice architectures.

Nevertheless, MiCSPARC is a robust method that enables the processing of microtubule specimens that may not meet traditional optimality criteria, thereby increasing reconstruction throughput. This may be particularly valuable for exploring nucleotide state transitions in motor proteins and managing heterogeneities in decorator binding modes. Factors like mask choice and alignment quality during signal subtraction can influence results, especially for varied decorators. To address this need for high throughput and ease-of-use, MiCSPARC also includes user-friendly GUI tools for visualizing angle smoothing accuracy and seam reconstruction, while also facilitating integration with CryoSPARC projects, as well as automated versions of the earlier steps of the pipeline. These tools have been made available to the academic community at: https://github.com/wieczoreklab/MiCSPARC.

The regulation of microtubule polymerization is closely linked to structural dynamics in α/β-tubulin [reviewed in (Brouhard and Rice, 2018)]. Yet, the field lacks a comprehensive atomic-level view of how structural transitions that take place in α/β-tubulin during various stages of GTP hydrolysis influence microtubule nucleation, growth, and disassembly. Additionally, local heterogeneities in microtubule lattice structure, particularly at the seam, are an intriguing target for structural investigations (Zhang et al., 2018), but these features have been difficult to dissect with existing cryo-EM SPA methods. Alternative computational pipelines like MiCSPARC, building on recent advances in SPA in cryo-EM, should provide a foundational suite of tools for dissecting these models at atomic resolution in the future.

## Acknowledgements

We thank Miroslav Peterek and Bilal Qureshi from ScopeM (ETH Zürich), as well as Pavel Afanasyev from the Cryo-EM Knowledge Hub (ETH Zürich), for cryo-EM sample preparation and data collection training and support on GDP microtubule specimens. We thank Mohamed Chami from BioEM Lab (University of Basel) for cryo-EM data collection on KIF5A-decorated GMPCPP microtubule specimens. We thank Christian Landolt, Alex Myczko, and Emil Zylis (ETH Zürich) for IT support. We thank Natacha Gaillard for providing the KIF5A motor domain sample. We are grateful to Stadt Zürich, Umwelt-und Gesundheitsschutz, and Fachbereich Veterinärdienste for supplying porcine brain tissue. We thank Sami Chaaban for testing early versions of MiCSPARC. This study includes calculations performed on the Euler cluster of ETH Zürich. This work was supported by startup funds from the ETH Zürich, SNSF Project Grants (#31003A_166608 to MOS, and #310030_208120 to MW), and an SNSF Starting Grant (#TMSGI3_211309 to MW).

## Author Contributions

DZ and MW designed the study. YX purified porcine tubulin, prepared undecorated GDP microtubule samples and grids, and collected cryo-EM data for undecorated GDP microtubules. KS initially processed data used to develop MiCSPARC with RELION, and SRC tested the MiCSPARC pipeline by comparing its performance with previous RELION pipelines. DZ developed MiCSPARC. DZ, HMH, PF and MW performed MiCSPARC-based cryo-EM processing and model building. HMH and PF implemented GUI and automated processing procedures and established the GitHub source for MiCSPARC. MW wrote the paper with contributions from all authors. MOS and MW supervised the study and obtained funding.

## Methods

### Protein purification

Calf brain tubulin used in the KIF5A motor domain-decorated dataset was purchased from the Centro de Investigaciones Biológicas Margarita Salas (Microtubule Stabilizing Agent Group), CSIC, Madrid, Spain.

Porcine brain tubulin used in the undecorated dataset was purified from brain tissue following the method of Castoldi and Popov (Castoldi and Popov, 2003).

Human kinesin KIF5A motor domain (residues 1–300, insert prepared from cDNA kindly provided by C. Hoogenraad) in PSTCm1 vector (Olieric et al., 2010) was expressed in Rosetta2 (Novagen) cells grown at 37 °C in LB medium supplemented with 50 μg/ml kanamycin and 30 μg/ml chloramphenicol to an OD600 of 0.4–0.6. Expression was induced with 0.4 mM isopropyl β-D-1-thiogalactopyranoside (IPTG; Sigma-Aldrich), and cultures were incubated overnight at 20 °C. Cell pellets were resuspended in lysis buffer (50 mM HEPES, pH 8.0, 500 mM NaCl, 10 mM imidazole, 10% glycerol, 2 mM β-mercaptoethanol, and one cOmplete EDTA-free protease inhibitor cocktail tablet (Roche)) and lysed on ice by ultrasonication. Lysates were clarified by ultracentrifugation, and the resulting supernatants were filtered through a 0.45 µm filter. The protein was affinity purified by immobilized metal affinity chromatography on a 5 ml HisTrap FF Crude column (GE Healthcare). The eluted fractions containing protein were pooled, concentrated and further purified by size-exclusion chromatography on a HiLoad 16/60 Superdex 200 column (GE Healthcare) equilibrated in 20 mM Tris-HCl, pH 7.5, 150 mM NaCl, 0.1 mM ADP, and 2 mM DTT. Eluted peak fractions were pooled, concentrated to 7.7 mg/ml and stored at −80 °C.

### Cryo-EM imaging of KIF5A motor domain-decorated, GMPCPP microtubules

To generate GMPCPP-stabilized microtubules, purified calf brain tubulin was resuspended at 4 mg/ml on ice using BRB80 buffer (80 mM PIPES, 1 mM EGTA, 1 mM MgCl2, pH 6.8) supplemented with 0.5 mM GMPCPP (Jena Bioscience). The tubulin solution was incubated on ice for 5 min then centrifuged for 10 min at 16,800 × g at 4°C. The supernatant was transferred to a fresh tube and placed on a 37°C heat block for 40 min. The GMPCPP-stabilized microtubule solution was snap-frozen in liquid nitrogen and stored at −80 °C until use.

To generate cryo-EM grid specimens containing KIF5A motor domain-decorated, GMPCPP-stabilized microtubules, freshly thawed KIF5A motor domain (7.7 mg/mL) was diluted with BRB80 to 2 mg/mL KIF5A, supplemented with 5 mM AMPPCP, and incubated for 30 min on ice. Meanwhile, GMPCPP-stabilized microtubules were thawed and warmed for 10 min at 37 °C, followed by centrifugation at 16,800 × *g* for 5 min at 25 °C. The supernatant was discarded and the microtubule pellet was resuspended in warm BRB80 buffer to a final tubulin concentration of 1.33 mg/mL. 3.5 µL of the stabilized microtubule solution was pipetted onto a glow-discharged Quantifoil grid (R 1.2/1.3, Cu, 200 mesh) and incubated for 60 s at room temperature. The grid was blotted by hand and 3 µL of the KIF5A-AMPCPP solution was added, the grid was transferred to a Vitrobot Mark IV (Thermo Fisher Scientific) at 100% humidity and 25 ^°^C, and incubated for a further 60 s. Grids were blotted for 1 s with a blot force of 2, plunge-frozen in liquid ethane, and stored in liquid nitrogen until data collection.

Data were acquired at the BioEM facility of the University of Basel on a Titan Krios electron microscope at 300 keV (Thermo Fisher), with a GIF Quantum LS Imaging filter (20 eV slit width) and a K2 Summit electron counting direct detection camera (Gatan). Movies were recorded at a magnification of 78,125X, corresponding to a pixel size of 0.64 Å. The total dose per movie was 65.0 e-/Å2, fractionated over 62 frames. Data collection statistics are also provided in Supplementary Table 1.

### Cryo-EM processing of KIF5A motor domain-decorated, GMPCPP microtubules

Data processing underwent the following steps, as outlined for the MiCSPARC pipeline.

Movies underwent beam-induced motion correction and CTF estimation by CryoSPARC’s patch-based methods.

A rough template for particle picking was generated using CryoSPARC’s filament tracer, with 360 Å filament diameter, and 82 Å separation distance between segments, corresponding to the heterodimeric repeat distance of α/β-tubulin. These were extracted and underwent 2D classification to select classes with a single clear tube in the 2D average, excluding picks with poor alignment, no, or multiple tubes.

To generate a more complete and more correctly assigned set of filament picks, these initial picks were used to guide an extrapolation algorithm to sample potential filament positions across the micrographs. To account for misassignments of filament ID, particularly in the case of densely packed or overlapping filaments, existing filament groups were first split at particles with an in-plane rotation greater than 10° from the preceding particle. Subsequently, a linear best fit was determined on a per-filament group basis, modelling the ideally straight arrangement of single particle microtubule preparations. In cases where this resulted in significant variation against more than 10% of the initial picks in the filament, extrapolation was instead performed using a quadratic best fit, modelling a smooth, continuous curve while minimizing chances of overfitting compared to higher order approximations. New extrapolated particle positions were again extracted and processed by 2D classification as previously described.

To create synthetic references from our data for protofilament sorting, a subset of particles were fed into six different helical refinement jobs with theoretical helical parameters corresponding to potential protofilament numbers in the dataset (protofilament number-start number: 11-3, 12-3, 13-3, 14-3, 15-4 and 16-4). The corresponding rise and twist for each helix is outlined in Appendix 1. After visual inspection of all helical refinement maps in ChimeraX, particles from the 14-3 map were selected for subsequent processing based on its well-resolved tubulins and KIF5A decoration. Particles were symmetry-expanded using theoretical helical parameters for a 14-3 microtubule, centered on a single protofilament, refined and subjected to signal subtraction. This centered protofilament reconstruction was then used to generate a series of volumes via MiCSPARC’s reference generation script (Appendix 1).

The resulting volumes were used as references for microtubule architecture sorting by one round of heterogeneous refinement. Similarly to MiRPv2 (Cook et al., 2020), class assignment within filament groups was “smoothened”, assigning each particle the modal value of the surrounding 7 particles. Stretches of filament where over 70% of the particles had been assigned a different class to the rest of the filament were reassigned into their own filament groups, while lower confidence differences in assignment were unified to match the rest of the microtubule. Particles corresponding to the 14-protofilament, 3-start microtubule, which made up 83% of the dataset, were selected for further refinement.

To perform rough alignment of the microtubule, the particles underwent helical reconstruction with non-uniform refinement using the theoretical helical parameters (rise 8.79 Å, twist −25.7°), global- and local-CTF refined, and finally reconstructed using helical refinement with no helical symmetry applied.

The angles in-plane of the micrograph (psi) and the microtubule axial plane (phi) were corrected alongside local refinements. Psi angles were first unified within filaments by quadratic fit through particles with angles within ±10° of the modal psi angle. To account for the somewhat random assignment of phi angles following helical refinement, the fit of 14 parallel linear models (corresponding to each protofilament) was optimized and the line of modal agreement was taken as the unifying value, ensuring that each particle was oriented such that each protofilament would be averaged in the same position as its neighbours. After refining the particles and performing an additional round of phi unification, the resulting particles were again refined and then symmetry expanded 14-fold using the refined helical parameters and subjected to a final local refinement without recentering.

Following rough alignment of the whole MT, a single protofilament was selected and masked using ChimeraX’s SEGGER tool (Pettersen et al., 2021). In particular, the protofilament with the most mixed population of decorator registers was selected, both to ensure that a tight mask did not cause any decorator signal to be removed during signal subtraction, and to provide adequate population sizes of both registers for register correction 3D classification.

The resulting volume and particles were shifted to be centered on a selected protofilament and the box size restricted to exclude the opposite side of the microtubule. Particles were signal subtracted to remove neighbouring protofilaments and locally refined to produce a roughly aligned single protofilament average, resulting in relatively high-resolution density for the tubulin heterodimers but poorer density for the decorator due to the combined occupancy of registers.

To separate the registers of the aligned protofilament, particles first underwent 3D classification with 10 classes in CryoSPARC’s “simple” initialization mode, with filter resolution set to 12 Å to force the classification to focus on the large decorator density. Classes with the strongest density of the KIF5A motor domain corresponding to particles of the same registers were locally refined together to produce two references of clear register difference.

These two references were used in a second round of 3D classification to separate the entire particle set into one of the two register classes. Each class was locally refined separately, and the class with worse reported resolution was chosen to be shifted by 41 Å (corresponding to a tubulin monomer) to align the register with the other class. The particles were finally combined, locally refined, and underwent duplicate removal, global and local CTF refinement to produce a final high-resolution average of a single register-corrected protofilament.

To produce a reconstruction of the whole microtubule, the separation of protofilament registers was restored using CryoSPARC’s heterogeneous reconstruction while retaining the refined per-particle CTF information.

To determine the seam position, protofilaments were grouped into sets produced from the same symmetry expanded particle, representing protofilaments of the same “layer” within the microtubule, and scored on the likelihood that they laid at the seam position of that layer based on the registers of the surrounding protofilaments. This process was repeated along each microtubule and averaged on a per-microtubule, producing a consensus seam position. The protofilament particles were un-symmetry-expanded by only retaining particles at the seam position.

Previous recentering and box size reduction to focus on single protofilaments were reverted to reconstruct a volume of the whole microtubule. Particles were locally refined and underwent duplicate removal followed by global and local CTF refinement to produce a final high-resolution average of the 14-3 seam-resolved KIF5A-decorated GMPCPP microtubule.

### Cryo-EM imaging of undecorated, dynamic microtubules

To generate spontaneously polymerized microtubules, 25 μM porcine brain tubulin in BRB80 buffer was supplemented with 1 mM GTP and 0.05% NP-40 (vol/vol) (Thermo Fisher Scientific) and was incubated at 37°C for 15 minutes. 3.5 μL of microtubule solution was then applied on glow discharged (25 mA, 30 s) C-flat™ holey thick carbon grid (R1.2/1.3, Cu, 300 mesh). After 30 s incubation in Vitrobot (FEI-Thermo Fisher) at 37°C, 100% humidity, the grid was blotted for 4 s using filter paper (Ted Pella Inc.) and plunge frozen in liquid ethane and stored in liquid nitrogen until data collection.

Movies were collected with a Gatan K3 in CDS mode and a slit width of 15-20 eV on a GIF BioQuantum energy filter on a TFS Titan Krios G3i (2) FEG of the ScopeM facility at ETH (Zürich, Switzerland). Automatic data collection was performed with “Faster acquisition mode” in EPU software (Thermo Fisher Scientific). The dataset was collected at 81,000 x magnification with a pixel size of 1.067 Å using counting mode. The total electron dose was 80 e-/Å2 over 40 frames with a defocus range varied between −2.8 and −1.2 μm.

### Cryo-EM processing of undecorated GDP microtubules

Data processing of undecorated microtubules proceeded in a similar fashion as outlined above for the decorated dataset, with some specific differences in parameters to adapt the process for the lack of an easily resolvable register marker.

Initial micrograph processing was performed as established. For the filament tracer, a filament diameter of 280 Å was used, adjusting to the smaller visual diameter of the microtubule due to the lack of a decorator protein. Particle picking and extrapolation, protofilament number sorting, and rough microtubule alignment otherwise proceeded as previously described.

During protofilament number sorting, a mixed population of 13 and 14-protofilament microtubules was found. These underwent rough alignment, symmetry expansion, and signal subtraction to expose single protofilaments separately, but protofilament particles from microtubules of both protofilament numbers were combined prior to register correction and refinement to allow for a consensus refinement of the highest resolution.

Due to the lack of a resolvable difference in register at lower resolutions, 3D classification for register correction proceeded with a filter resolution of 4 Å, allowing the difference in the density of the S9-S10 loop between α and β tubulin to be somewhat resolved and classified against. To aid this, a mask around a single central tubulin monomer was also used during classification.

Unlike in the decorated dataset, “ab initio” 3D classification only produced a strong average for one register of the protofilament. To circumvent this, a reference for the second register was created by recentering the volume by 41 Å using CryoSPARC’s volume alignment tool, and further 3D classification of the two protofilament registers proceeded as previously using these references.

Refinement of the single protofilament, and reconstruction of both 13 and 14-protofilament microtubules, proceeded as described for the KIF5A-decorated GMPCPP microtubule dataset.

### Model building

The KIF5A and microtubule models were generated based on an AlphaFold3 (Abramson et al., 2024) prediction using the sequences of human KIF5A (UniProt accession ID: Q12840), bovine tubulin α1B (P81947), bovine tubulin β2B (Q6B856), porcine tubulin α1A (P02550), and porcine tubulin β (P02554). The predicted structures were docked into their respective consensus maps using ChimeraX and subsequently refined manually in Coot (Emsley et al., 2010). Final real-space refinement was performed in PHENIX (Afonine et al., 2018). Model refinement statistics are provided in Supplementary Table S3.

## Supplementary Figures and Tables

**Supplementary Figure 1.**
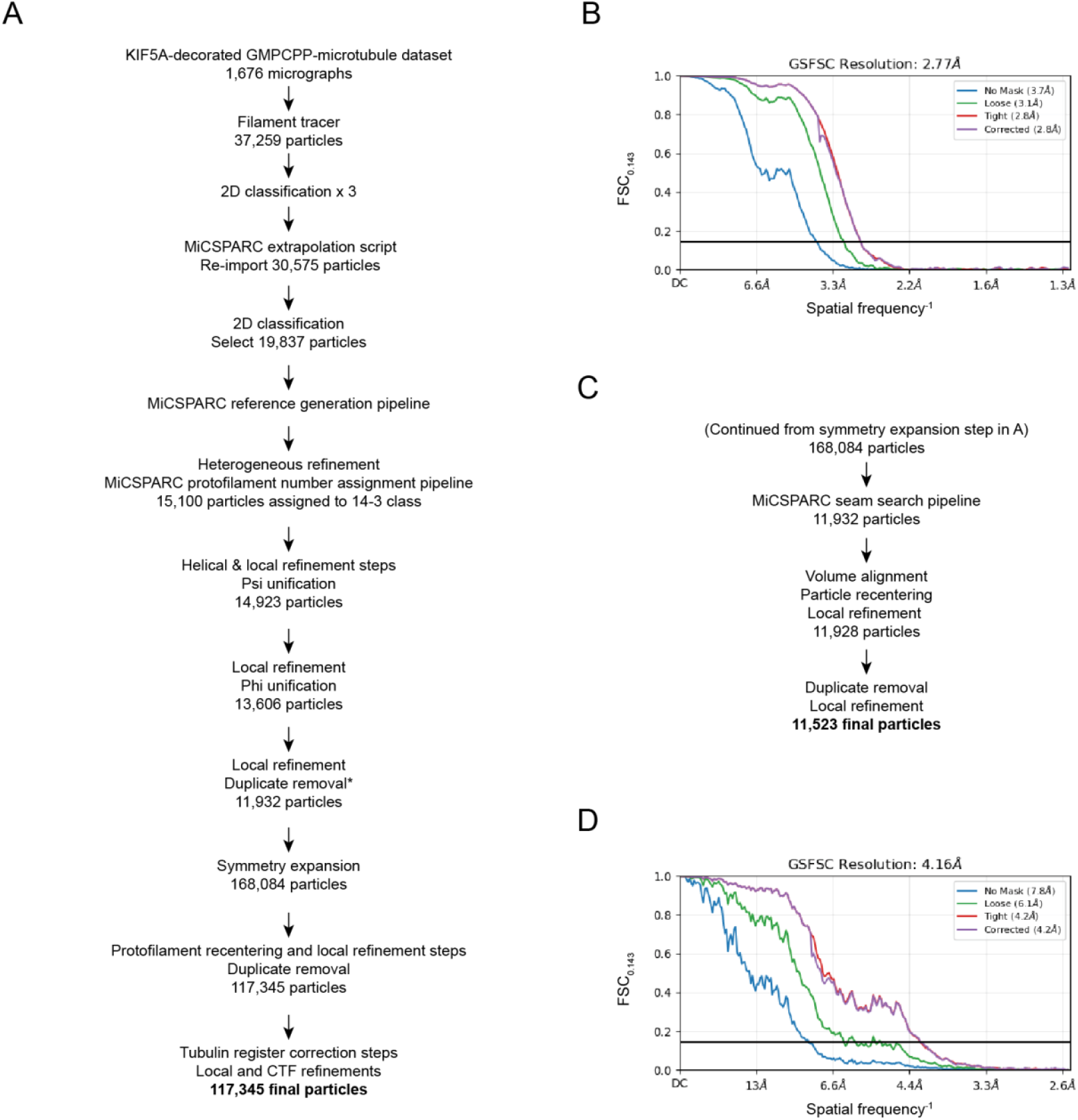
Data processing of KIF5A-decorated GMPCPP microtubules. **A)** Workflow for processing the tubulin register-corrected protofilament. **B)** Gold standard FSC curve for the protofilament reconstruction obtained in A). **C)** Workflow for processing the seam-corrected microtubule. **D)** Gold standard FSC curve for the seam-corrected microtubule reconstruction obtained in C).

**Supplementary Figure 2.**
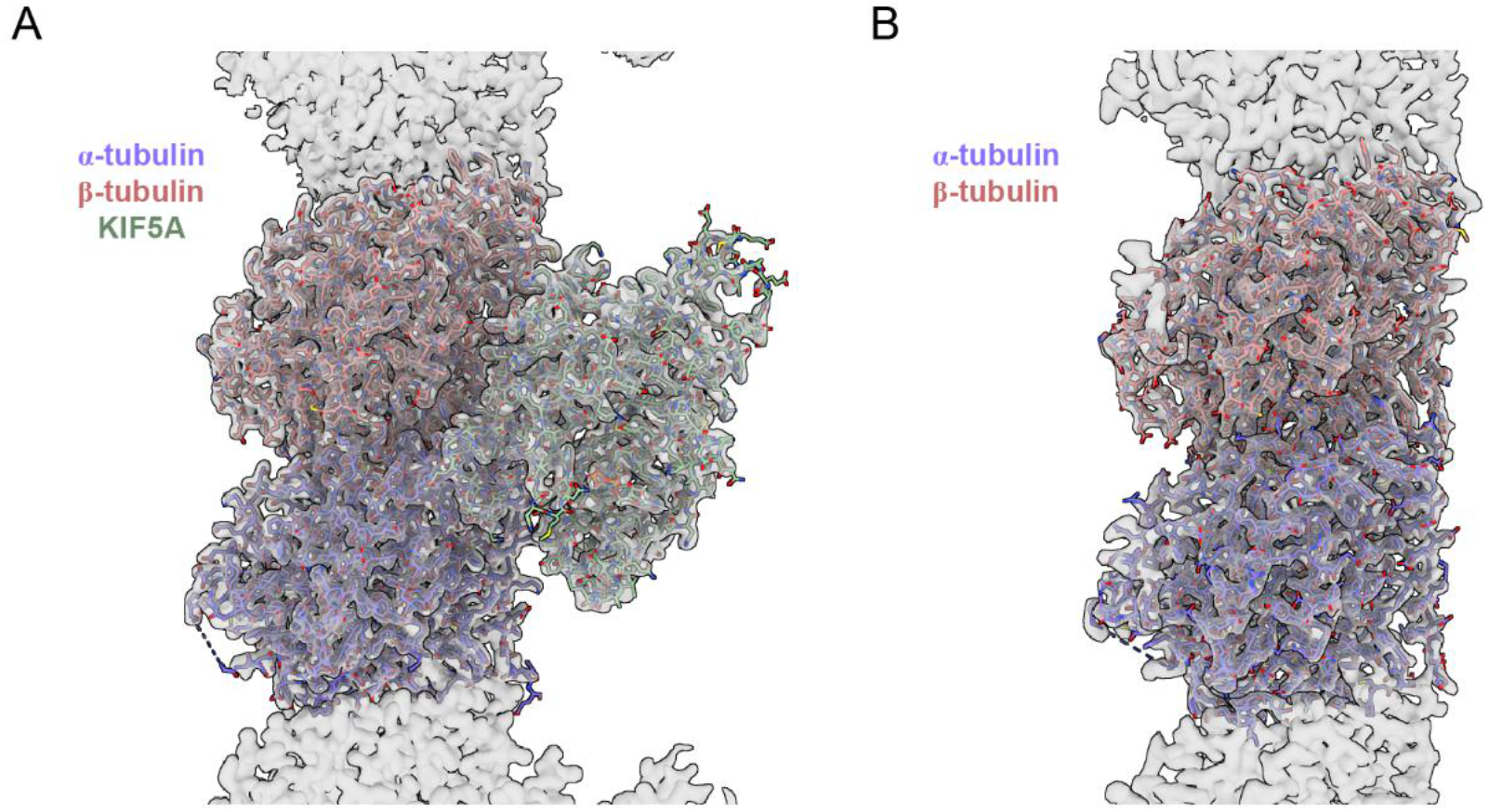
Stick representation views of KIF5A-decorated (**A**) and undecorated (**B**) α/β-tubulin models in the GMPCPP microtubule and GDP microtubule protofilament density maps (transparent grey surfaces). Maps were sharpened using EMReady (He et al., 2023).

**Supplementary Figure 3.**
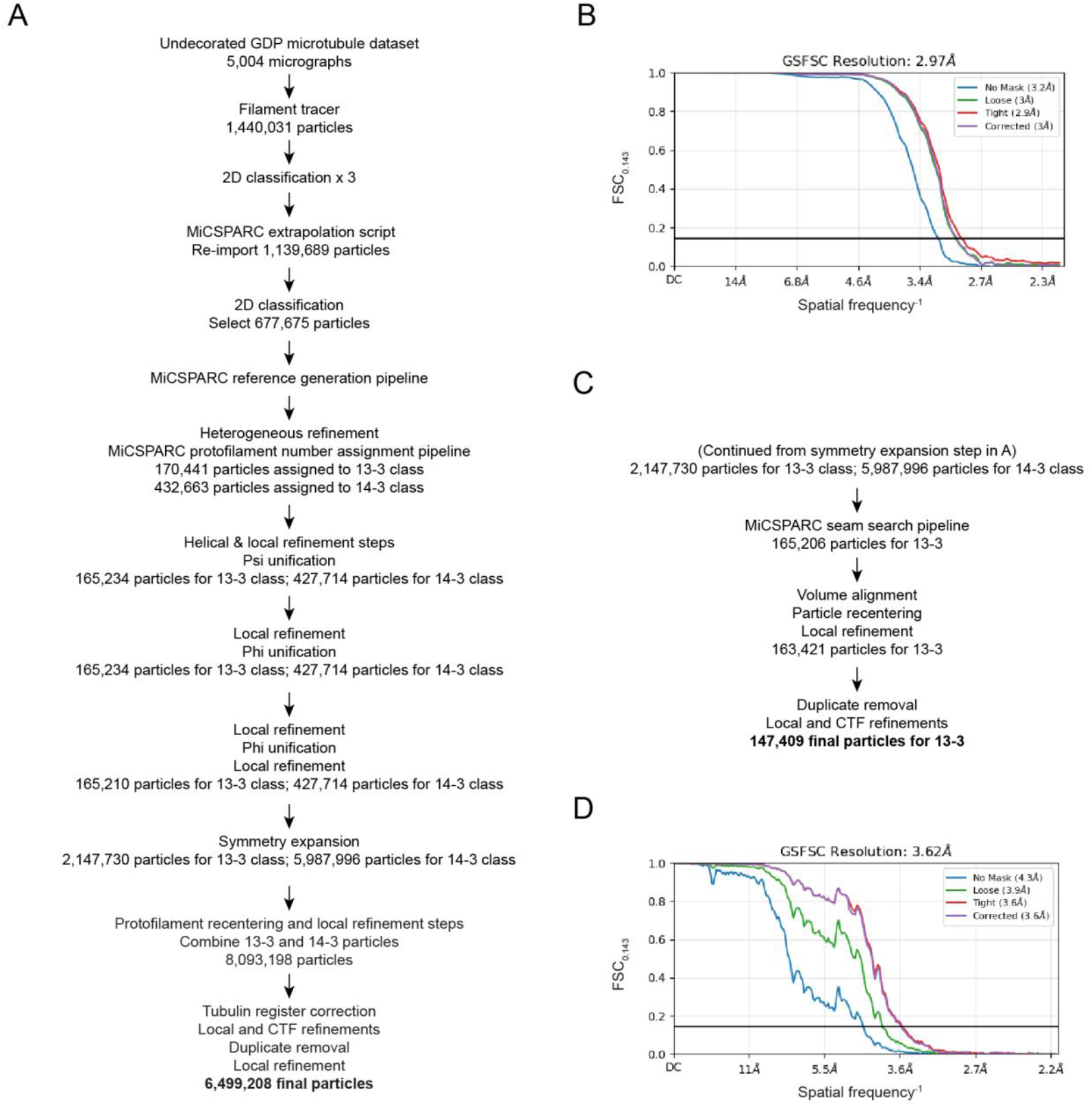
Data processing of undecorated GDP microtubules. **A)** Workflow for processing the tubulin register-corrected protofilament. **B)** Gold standard FSC curve for the protofilament reconstruction obtained in A). **C)** Workflow for processing the seam-corrected microtubule. **D)** Gold standard FSC curve for the seam-corrected microtubule reconstruction obtained in C).

**Supplementary Table S1.** Cryo-EM data collection.

|  | *KIF5A-decorated GMPCPP microtubules* | *Undecorated GDP microtubules* |
| --- | --- | --- |
| Voltage (keV) | 300 | 300 |
| Magnification | 78,125 X | 130,000 X |
| Pixel size (Å/px) | 0.64 | 1.06 |
| Camera | Gatan K2 | Gatan K2 |
| GIF slit width | 20 eV | 20 eV |
| Mode | Counting | CDS |
| Number of movies | 1,676 | 5,004 |
| Number of frames | 62 | 50 |
| Exposure time | 13 s | 3 s |
| Total dose (e/Å<sup>2</sup>) | 65 | 80 |

**Supplementary Table S2.** Cryo-EM data processing.

|  | *KIF5A-decorated*<br>*GMPCPP*<br>*protofilament* | *KIF5A-decorated, 14-*<br>*3 GMPCPP*<br>*microtubule* | *Undecorated*<br>*GDP*<br>*protofilament* | *Undecorated,*<br>*13-3 GDP*<br>*microtubule* |
| --- | --- | --- | --- | --- |
| Processing pixel size (Å/pixel) | 0.64 | 1.28 | 1.067 | 1.067 |
| Box size (pixels) | 512 | 512 | 256 | 512 |
| Symmetry imposed | C1 | C1 | C1 | C1 |
| Number of particles | 117,345 | 11,523 | 6,499,208 | 147,409 |
| Resolution (Å)<br>(FSC 0.143 cutoff) | 2.8 | 4.2 | 2.9 | 3.6 |
| Model building | Yes | No | Yes | No |
| Accession number | EMD-55425 | EMD-55444 | EMD-55431 | EMD-55440 |

**Supplementary Table S3.** Model building and refinement.

|  | *KIF5A-decorated GMPCPP microtubules* | *Undecorated GDP microtubules* |
| --- | --- | --- |
| Cross-correlation | 0.85 | 0.77 |
| Mean cross-correlation for ligands | 0.86 | 0.80 |
| Number of chains | 3 | 2 |
| Ligands | GTP: 1<br>Mg: 2<br>G2P: 1<br>ACP: 1 | GTP: 1<br>MG: 1<br>GDP: 1 |
| Number of atoms | 9,327 | 6,746 |
| Number of residues | 1,173 | 852 |
| MolProbity score | 1.91 | 2.02 |
| Clash score | 7.19 | 4.83 |
| Ramachandran outliers (%) | 0.69 | 0.59 |
| Allowed (%) | 2.67 | 4.62 |
| Favoured (%) | 96.65 | 94.79 |
| Bond length RMSD (Å) | 0.005 | 0.003 |
| Bond angles RMSD (°) | 0.665 | 0.685 |
| Accession number | PDB ID: 9T17 | PDB ID: 9T1D |

## Appendix 1: Suggested MiCSPARC pipeline

**CryoSPARC jobs** in bold blue

**MiCSPARC jobs** in bold orange

MiCSPARC commands highlighted in peach

ChimeraX commands highlighted in grey

### PARTICLE PICKING

#### Preprocessing

1. Run **Patch motion correction, Patch CTF estimation**, and **Curate exposures** jobs using default parameters

#### Generate initial templates for filament tracer

2 Run **Filament tracer**:
  ∘ ∼300 Å filament width*, 82 Å separation, 300-400 Å template-free diameter
  ∘ If the filament tracer is not picking any microtubules at this stage, increasing the standard deviation (SD) of the Gaussian blur (e.g. to 2.0) and/or increasing both hysteresis values in increments of 1 can help

3 **Inspect picks** if necessary, followed by **Extract from micrographs**:
  ∘ ∼550-600 Å box size*, Fourier crop (bin) particles to ∼4-5 Å/pixel
  ∘ If tube edges are close to the circular mask (especially with large decorators), expand the box size in later particle extractions

4 Run a **2D Classification** with default parameters
5 **Select 2D** classes with a single clear tube

#### Generate initial picks

6 **Filament tracer** using the templates selected in step 5
7 **Inspect picks** if necessary, followed by **Extract from micrographs**:
  ∘ ∼550-600 Å box size*, Fourier crop (bin) particles to ∼4-5 Å/pixel
8 **2D classification**:
  ∘ 50-100 classes, depending on the number of particles
  ∘ 2 final iterations
9 **Select 2D** classes with a clear tube to use for another **2D classification**:
  ∘ 50 classes
  ∘ 2 final iterations
  ∘ Disable sigma annealing by setting “Start annealing sigma” at iteration 200
10 **Select 2D** classes with a clear tube to use for another **2D classification**:
  ∘ 1 class
  ∘ **Export particles group** when finished

#### Extrapolate initial picks

The filament tracer will generally not pick along the entire microtubule, and may introduce errors in filament grouping. This can be rectified using MiCSPARC’s **Filament extrapolation** script:

11 Using the exported directory (it is easiest to run the scripts directly within the exported directory in order to be able to reimport the results group), run: `$ python /path/to/csparc_extrapolate_filaments.py −-i JX_particles_exported.cs …`
12 Run **Import result group** using the .csg result file from **Filament extrapolation**: (`/path/to/CS-project/exports/groups/JX_particles/JX_particles_extrapolated.csg`)
13 **Extract from micrographs**
  ∘ Same box size as in step 7
  ∘ Fourier crop (bin) particles to ∼2 Å/pixel
14 **2D classification**:
  ∘ 50-100 classes, depending on the number of particles
  ∘ 2 final iterations
15 **Select 2D** classes with a clear tube to use for another **2D classification**:
  ∘ 50 classes
  ∘ 2 final iterations
  ∘ Disable sigma annealing by setting “Start annealing sigma” at iteration 200

### PROTOFILAMENT NUMBER SORTING

#### Generate references

References are required for 3D classification of particles into groups of different protofilament numbers / microtubule lattice architectures. This classification tends to be very dependent on references with the correct factors such as decoration state, positioning, and heterogeneity, and potentially even expansion or compression of the microtubule lattice. Thus, it is highly recommended to create semi-synthetic references directly from the dataset, as follows:

16 Run one **Helical refinement** job for each type of microtubule lattice architecture potentially expected in the dataset
  ∘ Theoretical helical parameters are calculated in the form:

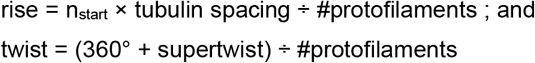
  ∘ Supertwist can generally be taken as 0 at this stage
  ∘ Common theoretical microtubule parameters (and hence **Helical refinements**):

17 Perform **Symmetry expansion** on the best helical refinement using its refined helical parameters
  ∘ Note: the “best” refinement is not necessarily the one with the highest resolution, but rather the one with the visually best-resolved protofilaments (i.e., no smearing around the tube, ideally distinct tubulin dimers)
18 Recenter the particles on a single protofilament by running **Volume alignment tools** with the “recenter to mask center of mass” option, but first:
  ∘ Create a protofilament mask in ChimeraX:
    ▪ First lowpass filter the best helical refinement to 15 Å using **Volume tools**
    ▪ Open map in ChimeraX, set an appropriate masking <u>threshold</u>
    ▪ Tools > Volume Data > Segment Map (may need to change “display at most” parameter to a number much greater than 60)
    ▪ Select a single protofilament
    ▪ File (in the Segment window) > Save selected regions to .mrc file (this is temporary, it doesn’t matter where they are saved)
    ▪ In the ChimeraX command line, run: `volume resample [new volume id] onGrid #1`
    ▪ Save the resampled volume and import into CryoSPARC using **Import volumes**
    ▪ Lowpass filter the protofilament volume to 15 Å using **Volume tools** (output as mask, and use the <u>threshold</u> chosen above)
19 **Downsample particles** to half of the current box size by cropping in real (not Fourier) space, and recenter particles
20 **Homogeneous reconstruction** of the downsampled particles (mask not required) followed by **Local refinement**:
  ∘ Use volume and particles from **Homogeneous reconstruction** as inputs, but use mask from step 18
  ∘ Use pose/shift gaussian prior during alignment with default values
  ∘ Allow recentering of shifts and rotations after each iteration
21 Use **Volume tools** to inverse the mask from step 20 and crop it in real space to half the box size
22 Perform **Particle subtraction** on the particles from step 20 using the mask from step 21
23 **Homogeneous reconstruction** of the subtracted particles (mask not required) followed by **Local refinement**:
  ∘ Use volume and particles from the **Homogeneous reconstruction** as inputs, but run **Volume tools** to generate and use a new, non-inverted mask from step 21
  ∘ Use pose/shift gaussian prior during alignment with default values
  ∘ Allow recentering of shifts and rotations after each iteration
  ∘ Enforce non-negativity
  ∘ At the end of the refinement, **Export volume group**
24 Run MiCSPARC’s **Reference generation** script: `$ python /path/to/csparc_create_pfn_references.py −-i JX_volume_exported.cs …` The `--recenter` coordinates are those reported by CryoSPARC in the volume alignment step 20 (“New center will be located at voxel coordinates:”). These coordinates must be input here in the format `--recenter “x, y, z”`

| Lattice architecture<br>(# pfs - helical “start”) | Rise (Å) | Twist (degrees) |
| --- | --- | --- |
| 11-3 | 11.2 | -32.7 |
| 12-3 | 10.3 | -30.0 |
| 13-3 | 9.46 | -27.7 |
| 14-3 | 8.79 | -25.7 |
| 15-4 | 10.9 | -24.0 |
| 16-4 | 10.3 | -22.5 |

#### Protofilament number assignment

25 Import the various reference volumes into CryoSPARC using **Import 3D volumes**
26 Run **Heterogeneous refinement** on the particles from step 15 using these imported volumes as references
  ∘ Force hard classification
  ∘ **Export particles group** when finished
27 Run MiCSPARC’s **Protofilament number assignment** script: `$ python /path/to/csparc_assign_pfns.py −-i …`
28 **Import result group** using the .csg result file from **Protofilament number assignment**
29 Run **Heterogeneous reconstruction** on the imported particles
  ∘ Note: an error will occur if there are any classes with zero particles from the step 26 heterogeneous refinement; in this case re-run the classification without that reference and repeat steps 27-29.
30 Run a **Split volume groups** job on the resulting particles

### ROUGH MICROTUBULE ALIGNMENT

Having each particle along the microtubule in the same orientation is important for later symmetry expansion and seam searching steps, as it allows us to assign each group of expansions as the same protofilament along the microtubule. For each good 3D class from steps 29-30 (or you can focus on just one dominant class, the number of particles is often enough):

31 Run a **Helical refinement** with the corresponding theoretical helical parameters
  ∘ Minimise per particle scale
  ∘ Use non-uniform refinement
  ∘ Note: particles may need to be unbinned and re-extracted if they hit Nyquist
  ∘ Note: if the “good” 3D class(es) contain a particle number greater than 50-100k, computational resources may be a problem due to the large box sizes required for unbinned microtubule segments combined with downstream symmetry expansion operations. In this case, it is probably best to repeat steps 1-30 with a randomized subset of micrographs (not particles, since particles must remain linked to filaments and micrographs in order for seam correction etc. to work).
32 Run a **Local CTF refinement** followed by a **Global CTF refinement**
  ∘ Use the particles, volume and mask from the **Helical refinement** as an input to the **Local CTF refinement**; then do the same for the **Global CTF refinement**, just update the particles to come from the **Local CTF refinement** jobo
33 Run a **Helical refinement** on the CTF-refined particles
  ∘ Do not impose helical symmetry (leave helical rise & twist fields blank)
  ∘ Minimise per-particle scale
  ∘ Use non-uniform refinement
  ∘ **Export particles group** when finished
34 Run MiCSPARC’s **Psi unification** script: `$ python /path/to/csparc_unify_psi.py …`
35 **Import result group** using the .csg result file from **Psi unification**
36 Run a **Local refinement**
  ∘ Use the resulting helical volume and mask from step 33
  ∘ Use pose/shift gaussian prior during alignment with 5 SD for rotations and 3 SD for shifts
  ∘ Allow rotation recentering
  ∘ **Export particles group** when finished
37 Run MiCSPARC’s **Phi unification** script: `$ python /path/to/csparc_unify_phi.py …`
38 **Import result group** using the .csg result file from **Phi unification**
39 Run a **Local refinement**
  ∘ Again use the resulting helical volume and mask from step 33
  ∘ Use pose/shift gaussian prior during alignment with 5 SD for rotations and 3 SD for shifts
  ∘ Allow rotation recentering
  ∘ (Optional) Force re-do GS split
  ∘ **Export particles group** when finished
40 Run MiCSPARC’s **Phi unification** script again: `$ python /path/to/csparc_unify_phi.py …`
41 **Import result group** using the .csg result file from the second **Phi unification**
42 Run a **Local refinement**
  ∘ Again use the resulting helical volume and mask from step 33
  ∘ Use pose/shift gaussian prior during alignment with 3 SD for rotations and 2 SD for shifts
  ∘ Do not allow any recentering
43 Perform **Symmetry expansion** on the refined particles
  ∘ Use refined helical parameters from step 33 with order = protofilament #
44 Run a **Local refinement**
  ∘ Again use the resulting helical volume and mask from step 33
  ∘ Use pose/shift gaussian prior during alignment with 3 SD for rotations and 2 SD for shifts
  ∘ Do not allow any recentering
  ∘ Note: for undecorated microtubules, if this step does not reach <4 Å resolution, later steps may prove difficult or even impossible. Decorated particles will be more lenient depending on the size of the decorator.

### PROTOFILAMENT ALIGNMENT

45 Create a mask around a single protofilament in the resulting volume from step 44 using ChimeraX, as in step 18
  ∘ Note: if refining multiple microtubule lattice arrangements, it can be helpful to select the protofilament that coincides best across all microtubule types.
  ∘ Note: with large decorators, select the protofilament with the most mixed population of decorator registers to ensure no gaps in the mask. Selecting the “worst” protofilament will also give the best split for the register classification, which will generally allow seam search to work better.
  ∘ Note: a tight mask along the protofilament and including the width of a single tubulin dimer plus the decorator can be beneficial at this stage; several masks may need to be tested to optimize the pipeline for different decorators.
46 Recenter the particles from step 44 on this protofilament by running **Volume alignment tools** with the “recenter to mask center of mass” option selected
47 Use **Volume tools** to crop the new realigned mask to half the original box size and invert
  ∘ Note: for this as well as the local refinement mask in step 49 below, it may be best to first re-run **Volume alignment tools** on the imported volume from step 45 and generate cropped masks on the centered volume. Generating masks from masks may not correctly propagate soft edges, thresholds etc.
48 **Extract from micrographs**
  ∘ Same box size as in step 47
  ∘ Recenter using all aligned shifts
49 **Homogeneous reconstruction** of the extracted particles (mask not required) followed by **Local refinement**:
  ∘ Use volume and particles from the **Homogeneous reconstruction** as inputs, but run **Volume tools** to generate and use a new, non-inverted mask from step 47
  ∘ Use pose/shift gaussian prior during alignment with 3 SD for rotations and 2 SD for shifts
  ∘ Do not allow any recentering
50 Perform **Particle subtraction** on the particles from step 49 using the mask from step 47
51 **Homogeneous reconstruction** of the subtracted particles (mask not required) followed by **Local refinement**:
  ∘ Use volume and particles from the **Homogeneous reconstruction** as inputs and use the non-inverted mask from step 49
  ∘ Use pose/shift gaussian prior during alignment with 3 SD for rotations and 2 SD for shifts
  ∘ Allow recentering of shifts and rotations after each iteration
  ∘ Enforce non-negativity

If refining multiple classes:

52 Pick a base refinement, align other protofilament number reconstructions with **Align 3D maps** (may need to be roughly aligned manually with volume alignment and ChimeraX first)
53 **Homogeneous reconstruction** of all combined particles (mask not required) followed by **Local refinement**:
  ∘ Use volume and particles from the **Homogeneous reconstruction** as inputs and use the non-inverted mask from step 49
  ∘ Use pose/shift gaussian prior during alignment with 3 SD for rotations and 2 SD for shifts
  ∘ Allow recentering of shifts and rotations after each iteration
  ∘ Enforce non-negativity

#### Protofilament register correction

54 Perform **3D classification** of the refined particles from step 53:
  ∘ 10 classes
  ∘ Filter resolution to 4 Å for undecorated microtubules and up to 12 Å for large decorators (lower resolution limits tends to allow the classification to work better for decorators)
  ∘ Per-particle scale = none
55 Identify “good” reference volumes:
  ∘ For undecorated microtubules: at least one class with a distinct α-tubulin S9-S10 loop, ideally at least two which exhibit a difference in register
  ∘ For decorated microtubules: at least one class with clear and convincing spacing of decorator, ideally at least two which exhibit a difference in register (if applicable)
56 For each “good” volume, perform **Homogeneous reconstruction** (mask not required) followed by **Local refinement**:
  ∘ Use volume and particles from the **Homogeneous reconstruction** as inputs and use the non-inverted mask from step 49
  ∘ Use pose/shift gaussian prior during alignment with 3 SD for rotations and 2 SD for shifts
  ∘ Allow recentering of shifts and rotations after each iteration
  ∘ Enforce non-negativity
57 If only one register was observed, create the other register using **Volume alignment tools**:
  ∘ Shift in Z by ∼41 Å
58 Perform **3D classification** of the refined particles from step 53
  ∘ 2 classes (use the refined volumes in step 56 as reference volumes)
  ∘ Filter resolution to 4 Å for undecorated microtubules and up to 12 Å for large decorators
  ∘ Input initialization mode
  ∘ Per-particle scale = none
  ∘ Note: for undecorated datasets, if everything is aligned well so far, this step should result in a ∼50/50 split of particles. The distribution of decorated datasets may depend on how well-resolved the seam was prior to symmetry expansion (step 43).
59 For each class, perform **Homogeneous reconstruction** (mask not required) followed by **Local refinement**:
  ∘ Use volume, particles and mask from the **Homogeneous reconstruction** as inputs
  ∘ Use pose/shift gaussian prior during alignment with 3 SD for rotations and 2 SD for shifts
  ∘ Allow recentering of shifts and rotations after each iteration
  ∘ Enforce non-negativity
60 Use **Volume alignment tools** to shift the lower-resolution class in Z by ∼41 Å
61 Use **Align 3D maps** to align the shifted class (maps & particles) to the unshifted, higher-resolution class
  ∘ Update particle alignments
62 Run **Homogeneous reconstruction** (mask not required) followed by **Local refinement**:
  ∘ Use volume, particles and mask from the **Homogeneous reconstruction** as inputs
  ∘ Use pose/shift gaussian prior during alignment with 3 SD for rotations and 2 SD for shifts
  ∘ Allow recentering of shifts and rotations after each iteration
  ∘ Enforce non-negativity
63 Run a **Local CTF refinement** followed by a **Global CTF refinement**
  ∘ Use the particles, volume and mask from step 62 as inputs to the **Local CTF refinement**; then do the same for the **Global CTF refinement**, just update the particles to come from the **Local CTF refinement** job
64 Run a **Local refinement**:
  ∘ Use volume and mask from the **Local refinement** in step 62, and particles from the **Global CTF refinement** in step 63, as inputs
  ∘ Use pose/shift gaussian prior during alignment with 3 SD for rotations and 2 SD for shifts
  ∘ Allow recentering of shifts and rotations after each iteration
  ∘ Enforce non-negativity

To further improve resolution of protofilament reconstruction:

65 Remove duplicate particles in the final refinement from step 64 using **Duplicate removal** with default settings
66 Perform **Reference-based motion refinement** on these particles using default settings
67 Run a **Local refinement**:
  ∘ Use volume, particles and mask from the **Local refinement** in step 62 as inputs
  ∘ Use pose/shift gaussian prior during alignment with 3 SD for rotations and 2 SD for shifts
  ∘ Allow recentering of shifts and rotations after each iteration
  ∘ Enforce non-negativity
  ∘ Sharpen the map either with **Sharpening tools** or with a deep learning method such as EMReady

### SEAM-CORRECTED MICROTUBULE RECONSTRUCTION

68 Perform a **Heterogeneous reconstruction** using pre-duplicate removal particles (step 64)
  ∘ Be sure to link the `alignments3D_multi` field of the 2-class 3D classification in step 58
  ∘ **Export particles group** when finished
69 Run MiCSPARC’s **Seam search** script: `$ python /path/to/csparc_seam_search.py …`
  ∘ Use the recenter coords from step 46, taking account for differences in the pixel size between step 46 and step 48 in case they were changed
70 **Import result group** using the .csg result file from the **Seam search** step (or groups, if analyzing multiple protofilament number microtubules; (e.g. `/path/to/CS-project/exports/groups/JX_particles/JX_particles_seamed_50_13pf.csg`)
71 Recenter the particles from step 68 on the coordinates output by the **Seam search** script using **Volume alignment tools**
72 **Extract from micrographs**
  ∘ Return to original microtubule filament box size
  ∘ Recenter using all aligned shifts
73 Run **Homogeneous reconstruction** (mask not required) followed by **Local refinement**:
  ∘ Use volume and mask from step 33, and particles from the **Homogeneous reconstruction** as inputs
  ∘ Use pose/shift gaussian prior during alignment with 3 SD for rotations and 2 SD for shifts
  ∘ Don’t allow any recentering
74 Run a **Local CTF refinement** followed by a **Global CTF refinement**
  ∘ Use the particles, volume and mask from step 73 as inputs to the **Local CTF refinement**; then do the same for the **Global CTF refinement**, just update the particles to come from the **Local CTF refinement** job
75 Run a **Local refinement**:
  ∘ Use volume and mask from the **Local refinement** in step 73, and particles from **Global CTF refinement** in step 74, as inputs
  ∘ Use pose/shift gaussian prior during alignment with 3 SD for rotations and 2 SD for shifts
  ∘ Don’t allow any recentering

Not strictly necessary, but to potentially improve the signal of decorators and/or the quality of the final microtubule reconstruction, one can try to:

76 Remove duplicate particles in the final refinement from step 75 using **Duplicate removal** with default settings
77 Perform a **3D Classification** job:
  ∘ 4-10 classes
  ∘ Filter resolution to 4 Å
  ∘ Force hard classification
78 Perform **Reference-based motion refinement** on particles from the best class using default settings
79 Run a **Local refinement** on these final particles:
  ∘ Use volume and mask from the **Local refinement** in step 75, and particles from step 78 as inputs
  ∘ Use pose/shift gaussian prior during alignment with 3 SD for rotations and 2 SD for shifts
  ∘ Don’t allow any recentering
80 Sharpen the map either with **Sharpening tools** or with a deep learning method such as EMReady

